# Gene expression in *Lucilia sericata* (Diptera: Calliphoridae) larvae exposed to *Pseudomonas aeruginosa* and *Acinetobacter baumanii* identifies shared and microbe-specific induction of immune genes

**DOI:** 10.1101/2021.04.02.438197

**Authors:** C.H. McKenna, D. Asgari, T.L. Crippen, L. Zheng, R.A. Sherman, J.K. Tomberlin, R.P. Meisel, A.M. Tarone

**Affiliations:** Department of Entomology, Texas A&M University, 2475 TAMU, College Station, TX 77843-2475; Department of Biology and Biochemistry, University of Houston; Southern Plains Agricultural Research Center, Agricultural Research Service, US Department of Agriculture, College Station, TX, 77845; Board Chair and Director, BioTherapeutics, Education & Research (BTER) Foundation, Irvine, California, USA; Monarch Labs, Irvine, California, USA

**Keywords:** maggot therapy, antibiotic resistance, wound debridement, RNA-seq, antimicrobial peptide

## Abstract

Antibiotic resistance is a continuing challenge in medicine. There are various strategies for expanding antibiotic therapeutic repertoires, including the use of blow flies. Their larvae exhibit strong antibiotic and antibiofilm properties that alter microbiome communities. One species, *Lucilia sericata*, is used to treat problematic wounds due to its debridement capabilities and its excretions and secretions that kill some pathogenic bacteria. There is much to be learned about how *L. sericata* interacts with microbiomes at the molecular level. To address this deficiency, gene expression was assessed after feeding exposure (1 hour or 4 hours) to two clinically problematic pathogens: *Pseudomonas aeruginosa* and *Acinetobacter baumanii*. The results identified immunity related genes that were differentially expressed when exposed to these pathogens, as well as non-immune genes possibly involved in gut responses to bacterial infection. There was a greater response to *P. aeruginosa* that increased over time, while few genes responded to *A. baumanii* exposure and expression was not time-dependent. The response to feeding on pathogens indicates a few common responses and features distinct to each pathogen, which is useful in improving wound debridement therapy and helps develop biomimetic alternatives.

## Introduction

*Acinetobacter* and *Pseudomonas* species are emerging as significant human pathogens throughout the world (Chopra et al., 2008). Naturally derived antibiotics, such as penicillin, are the primary tool used to treat bacterial infections, but many bacterial species have evolved resistance to antibiotics (CDC, 2020). Among thereatest concern are the ESKAPE pathogens (*Enterococcus faecium*, *Staphylococcus aureus*, *Klebsiella pneumoniae*, *Acinetobacter baumannii*, *Pseudomonas aeruginosa*, and *Enterobacter* spp.), which currently cause the majority of hospital infections and effectively “escape” the effects of antibacterial drugs (Boucher et al., 2009).

Blow flies (Diptera: Calliphoridae) are a promising natural system for identifying novel antibiotics (Pavillard and Wright, 1957; Sherman, 2014; Sherman et al., 2000, 2013). Blow flies colonize vertebrate remains, thus they are exposed to many potential pathogens (Anderson et al., 2001). Blow fly larvae engage in extra-corporeal digestion of decomposing resources that are often associated with diverse microbial communities (Weatherbee et al., 2017). Blow fly larvae excrete and secrete molecules with antibiotic, antibiofilm, and wound healing properties (Bexfield et al., 2008; Cazander et al., 2009; van der Plas et al., 2009). One consequence of this feeding strategy is that blow flies alter the bacterial communities associated with vertebrate carrion when they feed (Pechal et al., 2014; Tomberlin et al., 2017).

There are two potential approaches to take advantage of blow fly biology to treat open and infected wounds. The first is an organismal approach, where sterilized larvae are placed on a wound and allowed to feed until debridement is complete. This approach is currently FDA cleared (Sherman, 2014; Sherman et al., 2000, 2013), but requires cultural acceptance of insects as medical devices, which is unlikely to be uniform across individuals and cultures. This approach also requires a supplier to maintain colonies of sterile blow flies that will feed on dead, but not live, tissue–limiting use for this purpose (Sherman, 2014; Sherman et al., 2000). In the United States, only strains of *Lucilia sericata* Meigen (Dipter: Calliphoridae) are used for wound debridement (Sherman, 2014). The second approach is to develop biomimetic technologies based on blow fly biology (Tonk and Vilcinskas, 2017). To those ends, the molecular and immunological processes associated with blow fly feeding are dissected in order to develop treatments that mirror organismal processes without the need for maintaining and using actual blow flies. Both approaches to controlling infections would benefit from a dissection of the molecular processes involved in blow fly-bacterial interactions. Specifically, the efficacy of blow fly treatments can depend on which bacteria are present (Steenvoorde and Jukema, 2004; van der Plas et al., 2008) and understanding why could improve the utility of maggot therapy on non-responsive bacterial pathogens. For example, determining the mechanisms responsible for bacteria-specific blow fly responses could be used to improve blow fly strains through transgenic approaches (Linger et al., 2016).

A key component of the wound healing properties of blow fly larvae may be the antimicrobial peptides (AMPs) that they produce as part of their innate immune system (flies do not possess an adaptive immune system). The fly humoral innate immune response can be initiated by the immune deficiency (IMD) and Toll signaling pathways, ultimately activating families of AMPs and other effector proteins that kill bacteria (Hoffmann, 2003). Different pathogens activate different pathways and are targeted by different families of effectors resulting in at least three possible combinations between bacteria and fly immune systems: 1) matches, in which the fly immune system recognizes a bacterium and properly responds with a targeted antibacterial response; 2) the inability to recognize a pathogen by cellular receptors; or 3) the ability to recognize a pathogen, but the inability to express the proper antibacterial molecular components to kill that pathogen (Buchon et al., 2014; Hanson et al., 2019; Hoffmann, 2003). Much of what is known about fly immunity derives from research on *Drosophila* (Diptera: Drosophilidae), which are tuned to yeast laden foods and bacteria associated with decomposing plants. This distinction between the *Drosophila* and other flies is important as blow flies and their close relatives appear to express many more AMPs than *Drosophila*, suggesting a need for blow flies and their allied species to respond to a more diverse bacterial assemblages encountered on carrion or manure (Pöppel et al., 2015; Sackton et al., 2017). For example, the transcriptome of *L. sericata* contains at least 47 transcripts that encode putative antimicrbial peptides and other small effector proteins (Pöppel et al., 2015), whereas *Drosophila* have approximately 20 AMPs (Sackton et al., 2007). Lastly, fly infection studies usually evaluate infection initiated by septic wounding (Khalil et al., 2015; Pöppel et al., 2015; Sackton et al., 2017), instead of evaluating the feeding responses of flies, which is a major expected route of exposure to bacteria in natural environments (Tomberlin et al., 2017).

We aimed to address the natural immune response of *L. sericata* larvae to *P. aeruginosa* or *A. baumanii* (two ESKAPE pathogens) on an otherwise sterile food source by measuring gene expression in the larvae using RNA-seq. *P. aeruginosa* is regularly studied regarding its interactions with *L. sericata* (Andersen et al., 2010; Barnes et al., 2010; Cazander et al., 2009; Kerridge et al., 2005), while there is relatively little known about *A. baumanii* interactions with *L. sericata* (Hirsch et al., 2019; Vrancianu et al., 2020). Our approach allowed an assessment of the uniformity of fly immune responses to two different gram-negative human pathogens. It also allowed an assessment of whether the immune response consists primarily of induction of AMPs, other effectors, or other genes. Moreover, we subjected the larvae to both short (1 h) and long (4 h) exposures to bacteria to determine if the immune response is similar across time periods and bacterial treatments. These results identify both bacteria-specific and shared responses that the fly may use to regulate (or fail to regulate) *P. aeruginosa* and *A. baumanii*, which could inform the control of bacteria with blow fly products derived from organismal or biomimetic systems.

## Experimental Procedures

### Blow fly rearing and infection

*L. sericata* larvae were fed either *P. aeruginosa, A. baumannii*, or a control medium for 1 h or 4 h. The strains used were *P. aeruginosa* (Schroeter) Migula (ATCC^®^ 15692^™^; American Type Culture Collection, Manassas, Virginia) isolated from an infected wound and *A. baumannii* Bouvet and Grimont (ATCC^®^ 19606^™^) isolated from urine. The bacteria were prepared and stored at −80°C according to manufacturers’ instructions. The bacteria were cultured on Trypticase Soy Agar (TSA) 24 h in advance of experiments, and single colonies were isolated and passaged onto new TSA plates incubated 18–24 h at 37°C for use. The inoculum was brought to a 0.7 OD_600_ nm in phosphate buffered saline (PBS), enumerated by a tenfold serial dilution in PBS, spread onto TSA plates, and incubated 18–24 h at 37°C. This exposure concentration was selected based on a preliminary study performed to determine the highest concentration at which the larvae survived exposure to *P. aeruginosa* and *A. baumannii*.

The experiment was performed across 3 batches. Each batch consisted of four replicates per treatment. Each replicate consisted of a tube containing Medical Maggots^™^ larvae exposed to *P. aeruginosa, A. baumannii*, or PBS for 1 h or 4 h (24 tubes per batch). The larvae were from a single strain (maintained for 30-years) of *L. sericata* supplied by Monarch Labs (Irvine, CA, USA). Disinfected eggs were impregnated in sterile gauze and shipped overnight, with each batch arriving in a different shipment. Upon arrival, larvae were verified to be second instar, an approximate count was performed, and 5 larvae were placed in Trypticase Soy Broth (TSB) and incubated at 37°C overnight and monitored for bacterial growth to confirm sterility. Second instar larvae from each batch were aseptically removed from the shipping vial and aliquoted into each treatment tube upon arrival.

Larvae from each batch were exposed to 10 μL of 10^9^ cfu/ml of the bacterial treatment, *P. aeruginosa* (6.9 ± 1.7 x 10^9^ cfu/ml) or *A. baumannii* (8.6 ± 4.6 x 10^9^ cfu/ml), or control PBS added onto a slant of 2 mL of TSA within a 5 mL round bottom polypropylene tube (Thermo Fisher Scientific Inc., Wilmington, DE, USA) (Supplemental Figure S1). Eight replicate tubes were prepared for each bacterial treatment or control (24 tubes per batch). The tubes were incubated at room temperature for 30 min in a biosafety hood prior to use. Fifteen larvae were aseptically added to each tube, which was capped and incubated at room temperature for 1 or 4 h in a sterile environment.

After 1 h exposure, 10 larvae from 4 of each of the *P. aeruginosa*, *A. baumannii*, and control tubes were collected using sterile technique. This was repeated after 4 h exposure for larvae from the 4 remaining tubes of each treatment and control. The 10 larvae from each collection were placed into a 1.5 mL microcentrifuge tube (Thermo Fisher Scientific Inc., Wilmington, DE, USA) and immediately flash frozen in an isopropanol/dry ice mixture. Each of these tubes was stored at −80°C until subsequent RNA-seq library preparation.

### RNA-seq data collection

RNA-seq was used to measure gene expression in the 24 larval samples from each of the three batches of the experiment (72 samples total, Supplemental Table S1). For RNA extraction, five of the 10 larvae were removed from each 1.5 mL microcentrifuge tube and placed in a new 1.5 mL microcentrifuge tube. All tubes and consumables were RNAse-free. RNA was extracted in a two-step process. First, whole RNA was extracted using TriReagent (Sigma-Aldrich Corp., St. Louis, Missouri) preparation following manufacturer’s protocols. Briefly, five larvae were macerated in 1 mL of cold TriReagent. Following this, 50 mL of ice-cold BAN reagent (Molecular Research Center, Inc., Cincinnati, OH, USA) was added and the solution was vigorously mixed. Next, the tubes were centrifuged at 14,000 G at 4°C for 15 min to isolate the RNA from the DNA and proteins. Approximately 500 μL of the top clear layer was carefully removed with a pipet and added to 500 μL of ice-cold 100% isopropanol. The tubes were mixed via inversion three times and allowed to rest on ice for 10 min to precipitate the RNA. The precipitate was then centrifuged at 14,000 G at 4°C for 15 min. Next, the supernatant was completely removed, 1 mL of ice-cold 70% ethanol was added, the tube was centrifuged at 4°C for 5 min at 14,000 G, the ethanol supernatant was eluted, and any remaining ethanol was allowed to completely evaporate. The RNA was then dissolved in a 10μL mixture of 99 μL of DNase/RNase/Nucleotide-free water and 1 μL of SUPERase•In^™^ (Invitrogen, Life Technologies Incorporated, Grand Island, New York). Next, the RNA was further purified using a Qiagen RNeasy Micro Kit and on-column DNase treatment following manufacturer protocols (Qiagen Inc., Valencia, CA, USA). RNA was eluted into a 1:100μL mixture of SUPERase•In and DNase/RNase/Nucleotide-free water. RNA concentration and quality was assessed with a NanoDrop (Thermo Fisher Scientific Inc., Wilmington, Delaware) and an Agilent 2100 BioAnalyzer (Agilent Technologies Inc., Santa Clara, CA, USA). Each sample was stored at −80°C until RNA-seq library preparation.

Each of the 72 RNA samples was used to construct a separate paired-end RNA-seq libraries (Supplemental Table S1) using the TruSeq mRNA stranded kit. All 72 samples were sequenced on six separate flow cells (all samples were loaded onto all flow cells) on an Illumina Hi-Seq 2500 (Illumina, Inc., San Diego, California) with 125 bp paired-end reads following manufacturer protocols. All library preparation and sequencing were performed by the Genomics and Bioinformatics Services at Texas A&M AgriLife Research (College Station, TX, USA). Sequence cluster identification, quality prefiltering, base calling and uncertainty assessment were done in real time using Illumina’s HCS 2.2.68 and RTA 1.18.66.3 software with default parameter settings. Sequencer.bcl basecall files were demultiplexed and formatted into fastq files using bcl2fastq 2.17.1.14 script configureBclToFastq.pl. All RNA-seq data collected in this study are available from the NCBI Gene Expression Omnibus (accession GSE161305).

### RNA-seq data analysis

Multiple approaches were used to analyze the RNA-seq data because a published genome is not yet available for *L. sericata*. In one approach, RNA-seq reads were aligned to a previously assembled *L. sericata* transcriptome (Sze et al., 2012) along with previously identified transcripts of AMPs and other effectors from *L. sericata* (Pöppel et al., 2015). In the second approach, transcripts from the annotated reference genome of the closely related *Lucilia cuprina* (Anstead et al., 2015) served as the reference. Each reference has contrasting costs and benefits. The *L. sericata* transcriptome is from the same species where the RNA-seq data were collected, but it is an assembled transcriptome from RNA-seq data, not an annotation from a complete genome. It may therefore be missing genes that were not expressed in the sampled tissues, and it lacks full length transcripts for alternatively spliced genes. In contrast, the *L. cuprina* annotated genome overcomes the shortcomings associated with a transcriptome assembly, but it is from a different (yet closely related) species.

When the *L. sericata* transcriptome was used as the reference (Sze et al., 2012), reads were aligned to transcriptome “nodes” (from assembly 25_5) using kallisto v0.44.0 (Bray et al., 2016). A node can be considered to be one or more consecutive exons that are not alternatively spliced. We added to this reference transcriptome assembly a set of 47 previously annotated AMPs and other effectors from *L. sericata* (Pöppel et al., 2015). Next, redundancy between transcriptome nodes and annotated AMPs was identified using MegaBLAST (Morgulis et al., 2008) with the default parameters to search the transcriptome database with the AMPs as queries. We excluded 61 nodes with >95% DNA sequence identity to the set of 47 AMPs.

The output from the kallisto pseudoalignment of RNA-seq reads to *L. sericata* transcripts was used to test for differential expression across treatments (with or without bacteria) and time points (1 h and 4 h of exposure). Pseudocounts output by kallisto can be fractions because reads are assigned to transcripts using a probabilistic method (Bray et al., 2016). Pseudocounts were rounded to the nearest integer. Read counts were summed over all 6 lanes for each of the 72 treatment-time-batch combinations. Then a combination of the limma and edgeR packages in R were used to test for differential expression (McCarthy et al., 2012; Ritchie et al., 2015; Robinson et al., 2010). In all analyses, a gene was considered to be differentially expressed with a false discovery rate (FDR) less than 0.05 (Benjamini and Hochberg, 1995). FDR adjusted *P-*values (*P*_ADJ_) were calculated separately for each analysis. The specific linear models used in our analysis are described below. Fold-change language throughout the document reflects log_2_ fold-changes.

Three different approaches were taken when using the *L. cuprina* genome as the reference. In the first two, kallisto was used to align the RNA-seq reads to annotated transcripts from two different assemblies of the *L. cuprina* genome. The first assembly was deposited in NCBI GenBank in 2015 by the University of Melbourne (NCBI accession GCA_001187945.1). The second assembly was deposited in NCBI GenBank in 2017 by the i5k Initiative (NCBI accession GCA_000699065.2). The kallisto pseudocounts from alignment to annotated transcripts from each of these two assemblies were analyzed using limma and edgeR, as described above. In the third approach, TopHat v2.1.1 (Kim et al., 2013; Trapnell et al., 2009) was used to align the RNA-seq reads to the Melbourne assembly, with a mismatch rate and read-edit distance setting of four. We then tested for differential expression between control and treatment groups separately for each time point using Cuffdiff v2.2.1 (Trapnell et al., 2012, 2013). AMP genes and other effectors in the *L. cuprina* genome were retrieved from the genome annotations.

Two methods were used to identify genes that were differentially expressed between control and treatment samples when using limma/edgeR, both including batch as a random effect. Pairwise comparisons between control and exposed individuals at each time point (1 h or 4 h) were done by modeling the effect of whether the larvae were fed (i.e., exposed to) bacteria or the control media (*B*) on the expression of each gene (*E*), with experimental batch (*d*) as a random effect and residual (*e*) estimated across replicates: *E* ~ *B* + *d* + *e*. This analysis was performed separately for expression data from 1 h after exposure and data from 4 h after exposure. The RNA-seq data from *P. aeruginosa* and *A. baumannii* infection were analyzed separately.

A second type of analysis was performed using limma/edgeR to identify genes that are differentially expressed upon bacterial exposure by analyzing both time points simultaneously. In this approach, the expression of each gene depends on whether the larvae were fed bacteria or the control media, the length of time they fed (*T*), and experimental batch: *E* ~ *B* + *T* + *d* + *e*. We fit this model to the RNA-seq data from *P. aeruginosa* and *A. baumannii* separately. Amongst the genes that are differentially expressed upon exposure to a given bacteria, we tested for genes that are further affected by the length of exposure (1 h vs 4 h) by incorporating an interaction term between exposure and time: *E* ~ *B* + *T* + *B*×*T* + *d* + *e*. This analysis was only performed for genes with significant differential expression with respect to *B* because the interaction term (*B*×*T*) has no biological meaning for genes not responding to bacterial exposure (i.e., those with an insignificant *B* term).

## Results

### P. aeruginosa induces effector gene expression more than A. baumannii

Differentially expressed AMPs and other effectors were identified in *L. sericata* upon exposure to *P. aeruginosa* or *A. baumannii* for 1 h or 4 h (Supplemental Tables S2 and S3). In the first analysis, an assembled transcriptome of *L. sericata* served as the reference (Sze et al., 2012), with 47 annotated AMPs and other effectors added to it (Pöppel et al., 2015). Nine effector transcripts were upregulated (fold-change ranging from 1.94 to 3.27) in *L. sericata* upon exposure to *P. aeruginosa* (Figure 1). Four of the 9 transcripts have a significant interaction term between exposure and time (*P*_ADJ_ < 0.05), with expression that was upregulated more (fold-change of the effect of time and treatment on expression ranging from 1.66 to 2.76) upon longer exposure to *P. aeruginosa* (Figure 1). These included AMPs (*Lser-Dipt3* and *Lser-Dipt5*, which encode Diptericins), a non-AMP effector (*Lser-Edin7*, which is an *elevated during infection*, or *edin*, gene family member), and two transcripts encoding proline-rich peptides (*Lser-PRP1* and *Lser-PRP4*). Notably, *Lser-PRP4* was previously shown to be highly expressed in the larval crop of *L. sericata* upon infection with a mixture of *P. aeruginosa* and *S. aureus* (Pöppel et al., 2015). In contrast, only two AMPs were significantly up-regulated upon exposure to *A. baumannii* (*Lser-Atta3*, with a fold-change = 2.58, which encodes an Attacin, and *Lser-Dipt3*, fold-change = 2.08), and neither significantly increased expression with exposure time (Figure 2). Differential expression of other genes using this reference transcriptome was not done because the non-AMP transcripts are only assembled into fragments or nodes (Sze et al., 2012).

**Figure 1.**
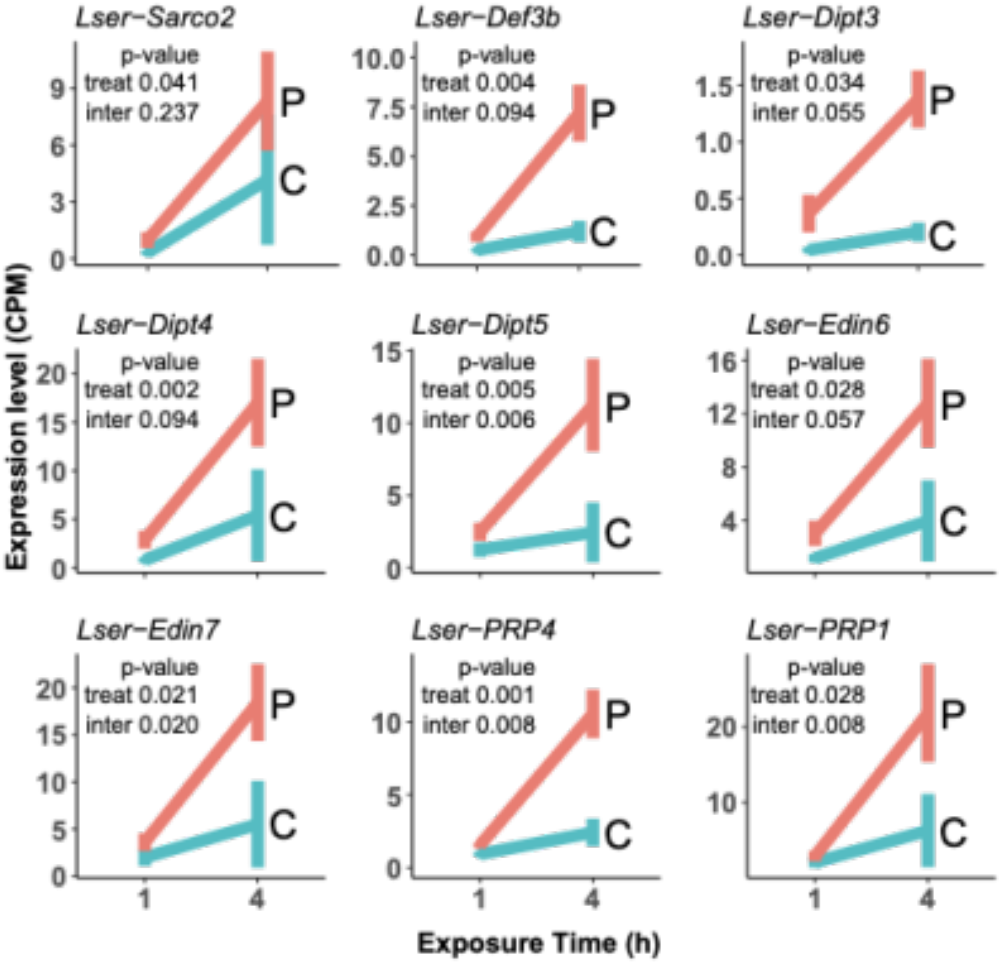
Transcripts encoding effector proteins that are upregulated upon exposure to *P. aeruginosa*. Gene expression was measured as counts per million (CPM). Salmon lines show expression when exposed to *P. aeruginosa* (P) and turquoise lines show expression in control samples (C). Expression was measured at 1 and 4 h. Text within each graph shows the FDR corrected *P*-value of a test for the effect of bacterial treatment on gene expression, and a test for the effect of an interaction between bacteria and time on expression (see Methods).

**Figure 2.**
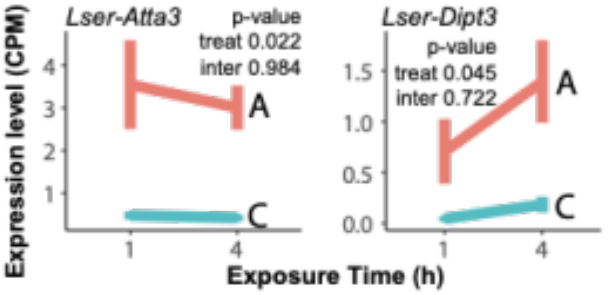
Transcripts encoding effector proteins that are upregulated upon exposure to *A. baumannii*. Gene expression was measured as counts per million (CPM). Salmon lines show expression when exposed to *A. baumannii* and turquoise lines show expression in control samples. Expression was measured at 1 hr and 4 h. Text within each graph shows the FDR corrected *P*-value of a test for the effect of bacterial treatment on gene expression, and a test for the effect of an interaction between bacteria on time on expression (see Methods).

### Differential expression of other genes

Next the annotated genome from the closely related *L. cuprina* (Anstead et al., 2015) was used as a reference to test for differential expression of AMP genes, non-AMP effector genes, other immune genes, and non-immunity genes upon exposure to *P. aeruginosa* or *A. baumannii* (Supplementary Tables S4, S5, S6 and S7). Two different genome assemblies (Melbourne and i5k) and two different analysis approaches were used to test for differential expression (limma/edgeR for both assemblies, and Cuffdiff for the Melbourne assembly). Because a reference genome from another species was used, only genes with a robust signal of differential expression were considered (i.e., they were detected as differentially expressed using all three combinations of reference genome and computational method) (Figure 3). Differential expression was tested separately for each length of exposure time (1 and 4 h). Note that it is not possible to directly compare AMP and other effector genes between the analyses using the *L. sericata* transcriptome and the different *L. cuprina* reference genomes because it is not feasible to assign direct orthology between species and reference genomes (due to the high evolutionary rate at which AMP genes are duplicated and the incomplete nature of genomic resources in these species). Therefore, the focus was on AMP and effector gene families common across analyses rather than orthologous genes between *L. cuprina* and *L. sericata*.

**Figure 3.**
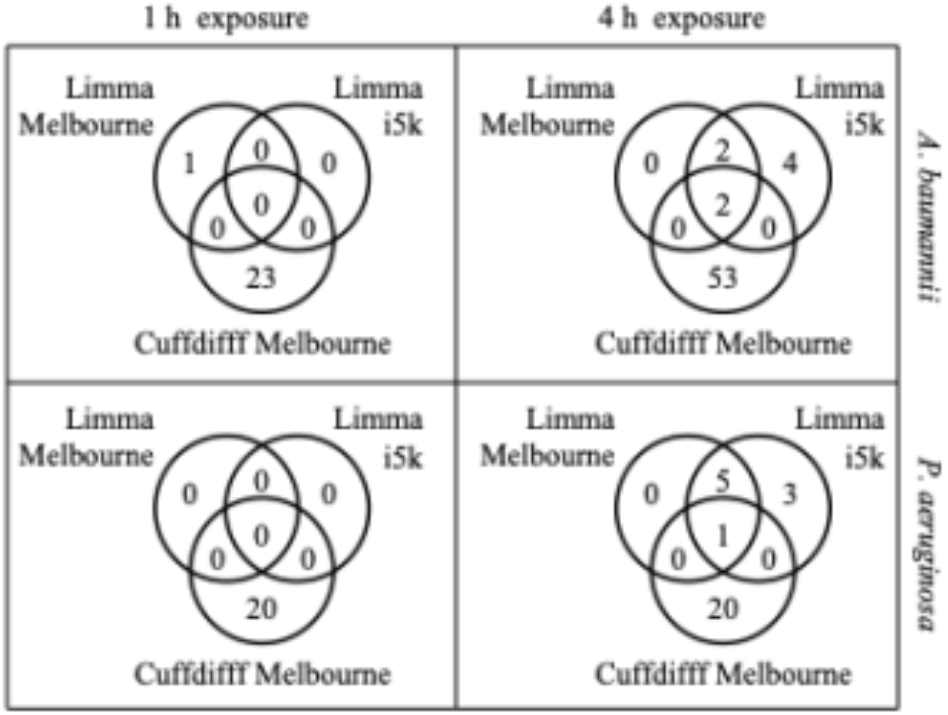
Venn diagrams of the number of differentially expressed genes detected using three different combinations of *L. cuprina* reference genomes (Melbourne and i5k) and analysis approaches (Limma and Cuffdiff). Complete lists of all genes are available in Supplemental Tables S4 and S5.

Differentially expressed genes in common across approaches were only detected for the 4 h bacterial treatments, and not the 1 h treatments. One gene (*FF38_08822*) was upregulated upon 4 h of exposure to *P. aeruginosa* (Figure 3, 4). This gene is homologous to *Drosophila melanogaster edin*, which encodes a short peptide that is induced by many bacteria (both Gram-negative and Gram-positive) and involved in the humoral immune response (Gordon et al., 2008; Vanha-Aho et al., 2012; Verleyen et al., 2006). Two *edin* transcripts were upregulated upon exposure to *P. aeruginosa* when using annotated *L. sericata* AMPs as a reference (Figure 1). Thus, the signal of upregulation of *edin* upon exposure to *P. aeruginosa* is robust to the reference transcriptome and analysis pipeline used.

**Figure 4.**
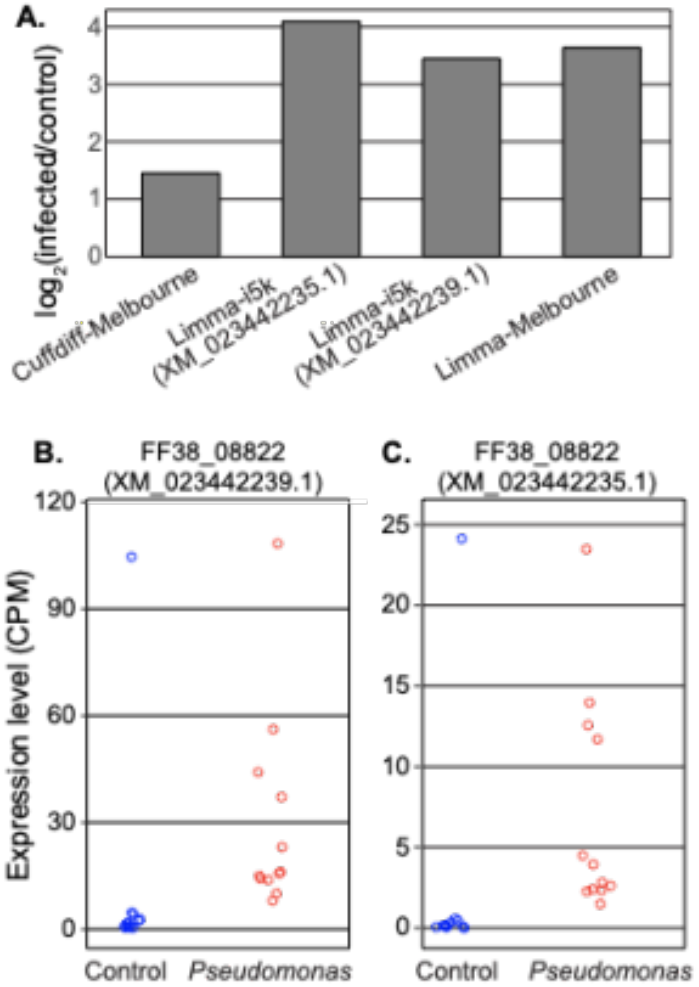
One gene was detected as differentially expressed upon exposure to *P. aeruginosa* at the 4 h time point in all three analyses using the *L. cuprina* reference genome (*FF38_08822*, homoogous to *edin*). Two different transcripts of the gene are present in the i5k reference genome (XM_023442239.1 and XM_023442235.1), and results are shown for each transcript separately. **A.** Log fold-change of differential expression between control and treatment for all three combinations of reference genomes (Melbourne and i5k) and analyses (Cuffdiff and Limma). **B-C.** Dots show the counts per million (CPM) values for each transcript of the gene in the control and *P. aeruginosa* treatment from each replicate. CPM values are taken from the Limma/i5k analysis.

One gene (*FF38_04326*) was upregulated and another (*FF38_01932*) was downregulated upon 4 h of exposure to *A. baumannii* (Figures 3, 5). *FF38_04326* encodes a protein that is homologous to *D. melanogaster* Pirk, a negative regulator of the Imd pathway (Kleino et al., 2008). The Imd pathway is triggered by DAP-type peptidoglycans found in the cell walls of Gram-negative bacteria (Kleino and Silverman, 2014), such as *A. baumannii*. *FF38_01932* is homologous to *D. melanogaster ecdysone-dependent gene 84A* (*Edg84A*), which is required for formation of the pupal cuticle in *D. melanogaster* (Apple and Fristrom, 1991).

**Figure 5.**
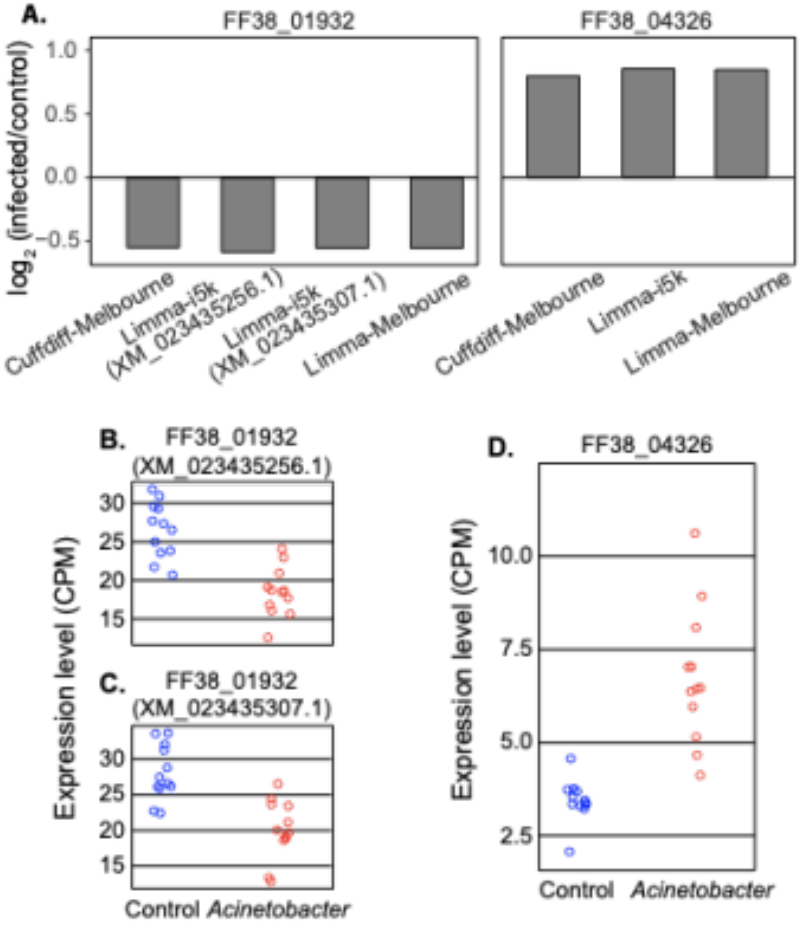
Two genes were detected as differentially expressed upon exposure to *A. baumanii* at the 4 h time point in all three analyses using the *L. cuprina* reference genome. *FF38_01932* is homologous to *D. melanogaster Edg84A*, and *FF38_04326* is homologous to *D. melanogaster pirk*. Two different transcripts of one gene are present in the i5k reference genome (XM_023435256.1 and XM_023435307.1), and results are shown for each transcript separately. **A.** Log fold-change of differential expression between control and treatment for all three combinations of reference genomes (Melbourne and i5k) and analyses (Cuffdiff and Limma) for each gene. **B-D.** Dots show the counts per million (CPM) values for each transcript of each gene in the control and *P. aeruginosa* treatment from each replicate. CPM values are taken from the Limma/i5k analysis.

There were 20–55 differentially expressed genes when evaluating the Melbourne reference genome with Cuffdiff, which greatly exceeds the number detected using limma/edgeR and either reference genome (Figure 3; Supplementary Tables S4, S5, S6, S7). Some genes distinct to the Cuffdiff analysis include three *Osiris* genes, a gene homologous to *D. melanogaster Peritrophin-15a/b*, *Tweedle* family genes *(Twdl)*, *Limostatin* (*Lst*), and *ecdysoneless* (*ecd*). The three *Osiris* genes (FF38_12518, FF38_12526, and FF38_12536, corresponding to *D. melanogaster Osi2*, *Osi9*, and *Osi15*, respectively) were downregulated (1.2, 1.6, and 4 fold, respectively) upon 1 h of exposure to *A. baumannii*. The *Osi15* homolog was also downregulated (4.3 fold) after 1 h of exposure to *P. aeruginosa* along with a 1.2 fold decrease in expression of a homolog of *Peritrophin-15a/b* (*FF38_03381*). *Osiris* genes, which are generally associated with chitin processes, are differentially expressed in immune responses, and similarly in the presence of a toxin, possibly through interactions with the peritrophic matrix (Smith et al., 2018). In the same 1 h time point, a *Twdl* gene (*FF38_01916*) was upregulated after exposure to both bacteria (1.1-1.3 fold). The *Lst* gene (*FF38_07809*) is upregulated 0.73 fold upon 4 h exposure to *P. aeruginosa*. Lst is a peptide hormone produced by endocrine corpora cardiaca cells during starvation, and it suppresses insulin production and secretion from insulin-producing cells by signaling through the G-protein coupled receptor PK1-R4 (Alfa et al., 2015). The *ecd* gene (*FF38_01073*) is upregulated 0.88 fold upon 4 h exposure to *A. baumannii*.

### AMPs are induced more than other genes

Finally, we used the *L. cuprina* genome to compare induction of AMPs with non-AMPs. The *L. cuprina* reference genome was used because the *L. sericata* transcriptome assembly can only be analyzed at the node level, not transcript / gene levels (Sze et al., 2012), which prohibits analysis of the expression to this end. The i5k *L. cuprina* genome served as a reference because it is the most up to date assembly. The limma/edgeR analysis was used to assess comparisons across time points. We analyzed the 1 and 4 h treatments together, as well as the 4 h treatment alone (no genes were differentially expressed at 1 h). Notably, this analysis is performed at the transcript-level. However, our comparison across *L. sericata* assemblies (Figure 3) was performed on genes because we could only determine correspondence across annotations at the level of genes. For this reason, the counts of differentially expressed transcripts do not necessarily correspond to the number of differentially expressed genes shown in the limma/i5k analysis reported in Figure 3.

AMP transcripts were more likely to be induced relative to other genes upon exposure to the bacteria. Six AMP genes were induced by exposure to *P. aeruginosa*, across both exposure times (i.e., 1 and 4 h), which encode Diptericin, Phormicin, Attacin, and Sarcotoxin proteins (Table 1). Only 6 non-AMP genes were induced by *P. aeruginosa*, which is a significantly smaller proportion than induced AMPs (Table 1). There are also 4 AMP genes induced upon exposure to *A. baumannii* across both exposure times, encoding Attacin, Diptericin, and Sarcotoxin proteins (Table 1). In comparison, there are only 3 non-AMP genes induced by *A. baumannii*, which is a significantly smaller fraction than the induced AMPs (Table 1). Of the genes upregulated after 4 h only, a similar excess of AMPs is observed (Table 1). We did not detect any AMP induction that depends on an interaction between exposure and time when using the *L. cuprina* reference (Supplemental Tables S6 and S7).

**Table 1.**
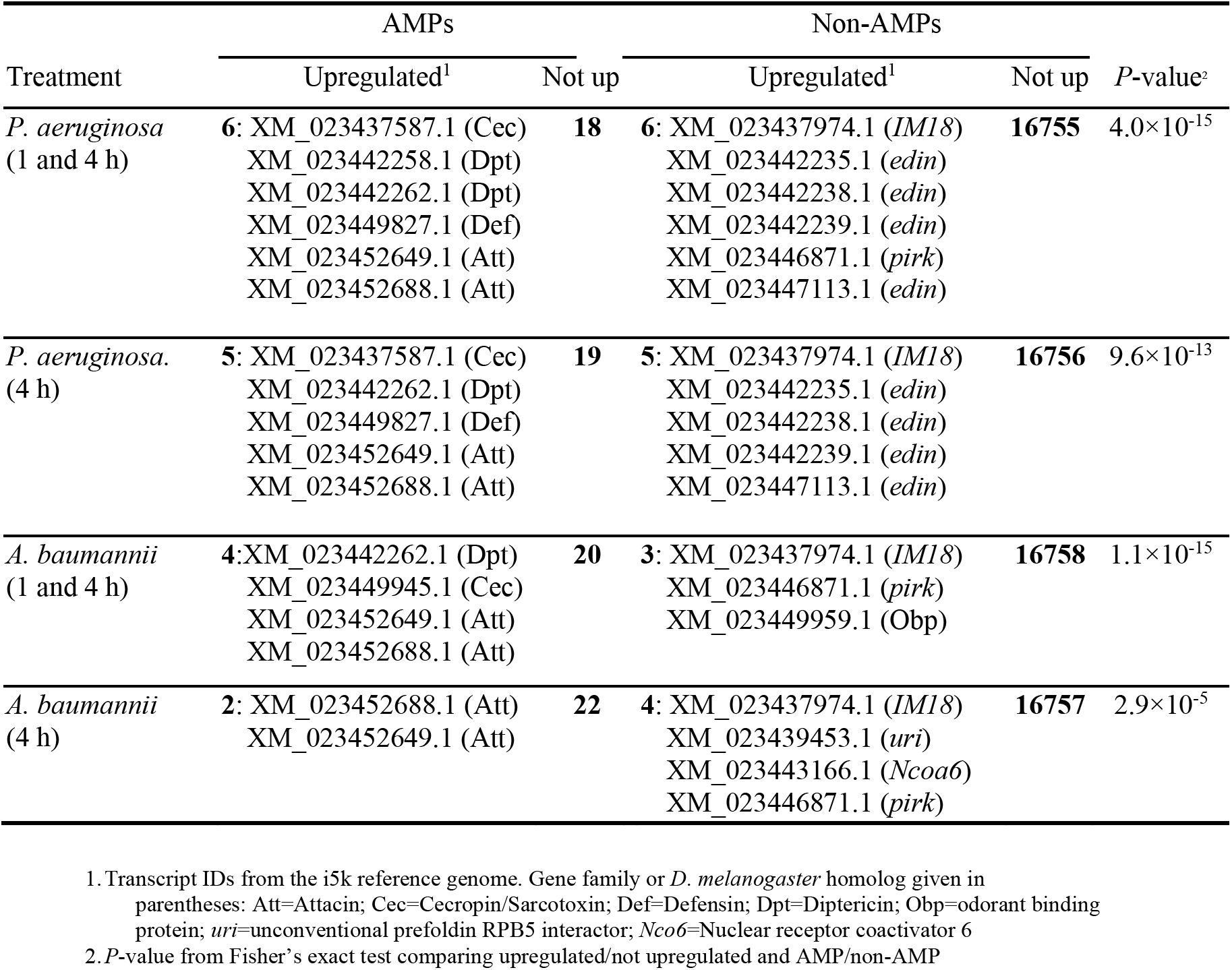
Induction of AMPs and non-AMPs.

| Treatment | AMPs |  | Non-AMPs |  | *P*-value<sup>2</sup> |
| --- | --- | --- | --- | --- | --- |
|  | Upregulated<sup>1</sup> | Not up | Upregulated<sup>1</sup> | Not up |  |
| *P. aeruginosa*<br>(1 and 4 h) | **6:** XM_023437587.1 (Cec)<br>XM_023442258.1 (Dpt)<br>XM_023442262.1 (Dpt)<br>XM_023449827.1 (Def)<br>XM_023452649.1 (Att)<br>XM_023452688.1 (Att) | **18** | **6:** XM_023437974.1 (*IM18*)<br>XM_023442235.1 (*edin*)<br>XM_023442238.1 (*edin*)<br>XM_023442239.1 (*edin*)<br>XM_023446871.1 (*pirk*)<br>XM_023447113.1 (*edin*) | **16755** | 4.0×10<sup>-15</sup> |
| *P. aeruginosa*<br>(4 h) | **5:** XM_023437587.1 (Cec)<br>XM_023442262.1 (Dpt)<br>XM_023449827.1 (Def)<br>XM_023452649.1 (Att)<br>XM_023452688.1 (Att) | **19** | **5:** XM_023437974.1 (*IM18*)<br>XM_023442235.1 (*edin*)<br>XM_023442238.1 (*edin*)<br>XM_023442239.1 (*edin*)<br>XM_023447113.1 (*edin*) | **16756** | 9.6×10<sup>-13</sup> |
| *A. baumannii*<br>(1 and 4 h) | **4:** XM_023442262.1 (Dpt)<br>XM_023449945.1 (Cec)<br>XM_023452649.1 (Att)<br>XM_023452688.1 (Att) | **20** | **3:** XM_023437974.1 (*IM18*)<br>XM_023446871.1 (*pirk*)<br>XM_023449959.1 (Obp) | **16758** | 1.1×10<sup>-15</sup> |
| *A. baumannii*<br>(4 h) | **2:** XM_023452688.1 (Att)<br>XM_023452649.1 (Att) | **22** | **4:** XM_023437974.1 (*IM18*)<br>XM_023439453.1 (*uri*)<br>XM_023443166.1 (*Ncoa6*)<br>XM_023446871.1 (*pirk*) | **16757** | 2.9×10<sup>-5</sup> |

1. Transcript IDs from the i5k reference genome. Gene family or *D. melanogaster* homolog given in parentheses: Att=Attacin; Cec=Cecropin/Sarcotoxin; Def=Defensin; Dpt=Diptericin; Obp=odorant binding protein; *uri*=unconventional prefoldin RPB5 interactor; *Ncoa6*=Nuclear receptor coactivator 6 2. *P*-value from Fisher's exact test comparing upregulated/not upregulated and AMP/non-AMP

Most of the non-AMP genes that are upregulated upon bacterial exposure when using the *L. cuprina* reference genome also encode immune genes. This includes four *edin* genes that are upregulated upon *P. aeruginosa* feeding (Table 1), similar to what was detected when using the *L. sericata* transcriptome as a reference (Figure 1). Upregulation of a *pirk* homolog was observed after exposure to *P. aeruginosa* or *A. baumannii* (Table 1), which was also detected in *A. baumannii* fed flies across our three different analysis approaches (Figure 5). Another gene responding to both bacteria is homologous to *D. melanogaster Immune induced molecule 18* (*IM18*). This gene was not annotated in the *L. cuprina* reference genome because it has low nucleic and amino acid identity to its homologs in other flies. *IM18* homologs have been identified as expressed during the immune response to bacteria across flies, consistent with being a non-AMP immune effector (Pei et al., 2014; Uttenweiler-Joseph et al., 1998).

Three genes with no direct relationship to immune response were also upregulated upon treatment with *A. baumannii*. First, a gene encoding an odorant binding protein (OBP) was upregulated when the 1 and 4 h *A. baumannii* data were considered together (Table 1). Many insect OBPs perform functions outside of olfaction (Pelosi et al., 2018; Sun et al., 2018), including mediating host-symbiont interactions (Benoit et al., 2017). OBP genes have also been identified as upregulated upon bacterial infection in *D. melanogaster* (Levy et al., 2004), and OBPs are components of the vertebrate humoral innate immune response (Bianchi et al., 2019). Second, a gene homologous to *D. melanogaster unconventional prefoldin RPB5 interactor* (*uri*) was upregulated after exposure to *A. baumannii* for 4 h (Table 1). In *Drosophila*, *uri* encodes a molecular chaperone involved in the transcriptional response to nutrient signaling and maintenance of DNA integrity via its interaction with Type 1 protein phosphatase (Kirchner et al., 2008). Uri is also implicated in the regulation of cell death, in part via TOR signaling (Chaves-Pérez et al., 2018). These results suggest that differential expression of *uri* might be involved in regulating the immune response to bacterial infection, as is indicated by others (Chaves-Pérez et al., 2018; Lynham and Houry, 2018). Third, a gene homologous to *D. melanogaster Nuclear receptor coactivator 6* (*Ncoa6*) was upregulated after exposure to *A. baumannii* for 4 h (Table 1). Nuclear receptors are an important component of intracellular signaling during some immune responses (Glass and Ogawa, 2006). *Ncoa6* encodes a protein that is part of a complex essential for regulating the transcription of genes involved in tissue growth via the Hippo signaling pathway (Qing et al., 2014). Hippo signaling is involved in the *Drosophila* innate immune response (Hong et al., 2018; Liu et al., 2016). Yorkie, a transcription factor regulated by the Hippo pathway, suppresses the Imd and Toll pathways, which are the two primary pathways responsible for inducing AMPs (Dubey and Tapadia, 2018). This provides a functional link between *Ncoa6* expression and response to bacterial exposure.

No AMP transcripts were detected as downregulated using the analysis that produced Table 1, but 3 non-AMP transcripts are downregulated in the *A. baumanii* 4 h treatment (Supplementary Table 5, Supplementary Figure 2). Two of these transcripts (*XM_023435256* and *XM_023435307*) are predicted to encode cuticle proteins that contain chitin binding domains (homologous to *D. melanogaster Ccp84A* genes). The third downregulated transcript (*XR_002762756*) is predicted to be a small nucleolar RNA that functions in the site-specific 2’-O-methylation of 18S RNA U-1356 (Galardi et al., 2002; Huang et al., 2005).

## Discussion

RNA-seq was used to identify genes that are differentially expressed when second instar *L. sericata* larvae feed on either *P. aeruginosa* or *A. baumannii* for 1 h or 4 h. The expression response included expected immunity genes, though the number of differentially expressed genes was not as great as seen with assays in which calyptrate flies were directly injected with microbes (Sackton et al., 2017). Specifically, the most sensitive analysis of our data revealed a maximum of 55 differentially expressed genes in *L. sericata* exposed to *A. baumannii* for 4 h (Figure 3). In contrast, adult house flies (*Musca domestica*) injected with a cocktail of *Serratia marcescens* and *Enterococcus faecalis*, differentially expressed 1,675 genes (784 upregulated and 891 downregulated), which represents >11% of all house fly genes (Sackton et al., 2017). In *L. sericata*, AMP and other effector genes are more likely to be induced than non-effector genes regardless of the bacteria, but exposure to *A. baumanii* also leads to the upregulation of non-effector genes (Table 1; Figure 3). In addition, more AMPs are upregulated by *P. aeruginosa* than *A. baumannii*, and only *P. aeruginosa* induction increases with the time of exposure (Figure 1; Figure 2).

### Immune genes are upregulated by feeding on bacteria

Both the *L. sericata* transcriptome and *L. cuprina* genome were used as a reference to identify upregulation of AMPs and other effector genes. We cannot directly compare orthologous genes between reference genomes due to the dynamic evolution of AMPs and other effectors (Sackton et al., 2007, 2017). Instead, we identified common gene families that are induced regardless of the analysis approach, which is suggestive of a robust signal of induction. The AMPs with such a robust signal of induction are Diptericins (*Lser-Dipt3*, *Lser-Dipt4*, and *Lser-Dipt5* in the *L. sericata* transcriptome), Cecropins/Sarcotoxins (*Lser-Sarco2*), Defensins (*Lser-Def3b*), Attacins (*Lser-Atta3*), and Edins (*Lser-Edin6* and *Lser-Edin7*). Moreover, two genes encoding proline-rich peptides (PRPs) are also induced by *P. aeruginosa* (Figure 1); these PRPs are not annotated in the *L. cuprina* reference because it is difficult to identify homologies of PRP genes across species (Scocchi et al., 2011; Yi et al., 2014). All AMP families identified are active against Gram-negative bacteria (Bulet et al., 1999; Hoffmann, 2003), such as *P. aeruginosa* and *A. baumanni*. Therefore, these AMPs appear to be important components of the immune response of *L. sericata* larvae to Gram-negative bacteria in their environment.

Diptericins may play a common role in response to gram negative bacteria in *L. sericata*. One of the nine *L. sericata* AMP transcripts upregulated upon exposure to *P. aeruginosa* (*Lser-Dipt3*) was shared with *A. baumannii* (Figure 1, Figure 2). *Diptericin* genes were also upregulated by *P. aeruginosa* and *A. baumannii* in the *L. cuprina* guided analysis (Table 1). Diptericins can confer resistance to infection by Gram-negative bacteria in *Drosophila* (Tzou et al., 2002). Our results suggest that Diptericins are a primary AMP induced when *L. sericata* is exposed to Gram-negative bacteria.

A homolog of *pirk* (*FF38_04326*) was upregulated upon exposure to both bacteria (Figure 3; Table 1). Pirk is a negative regulator of the Imd pathway, and it is activated shortly after infection in *Drosophila* (Kleino et al., 2008; Kleino and Silverman, 2014). Imd is involved in stimulating AMP gene expression in response to Gram-negative bacteria (Kleino and Silverman, 2014). Pirk functions to suppress the over-stimulation of the Imd pathway and over-expression of AMPs.

A poorly characterized effector, *IM18*, was in the list of “non-AMPs” commonly upregulated by both bacteria (Table 1). *IM18* genes exhibit low sequence identities across flies. However, protein similarities establish the homology of *L. cuprina IM18* to genes found in flesh flies, house flies, and *Drosophila* (e.g., 75% aa identity with *DOY81_008491* in *Sarcophaga bullata*; and 45% aa identity with *IM18* in *Drosophila virilis*, mostly at the C terminal end). *IM18* encodes a short peptide (~63 aa long in *L. cuprina* and 71 aa in *D. melanogaster*), which is in the same size class as known AMPs (Imler and Bulet, 2005; Lemaitre and Hoffmann, 2007). The *L. cuprina* peptide contains a prokaryotic membrane lipoprotein lipid attachment site profile in the first 19 aa (Sigrist et al., 2010), which implies direct interaction with bacterial cells. In *D. melanogaster* there is evidence that *IM18* could encode a non-canonical AMP (Pei et al., 2014; Uttenweiler-Joseph et al., 1998), which may be interesting as it seems to evolve differently than other AMP genes (Sackton et al., 2007); *IM18* has remained single/low copy but is divergent at the sequence level across species. Induction of *IM18* in these experiments is somewhat surprising because it is thought to be regulated by Toll (Seto and Tamura, 2013; Uttenweiler-Joseph et al., 1998). Activation of the Toll pathway was unexpected because flies were fed Gramnegative bacteria, and Toll signaling is generally associated with Gram-positive bacteria (Michel et al., 2001). However, there is some evidence from *D. melanogaster* that Gram-negative bacteria can also induce *IM18* expression (Short and Lazzaro, 2013), and other evidence suggests complex relationships between Imd and Toll responses (Nishide et al., 2019; Valanne et al., 2011). More research on the evolution, regulation, and function of *IM18* appears to be warranted given its distinct features compared to canonical AMPs and the shared response of this gene in *L. sericata* to two different ESKAPE bacteria.

Several effectors were distinct to the *L. sericata* response to *P. aeruginosa*, and not induced upon exposure to *A. baumannii*. Transcripts encoding Edins (*Lser-Edin6* and *Lser-Edin7*), a Defensin (*Lser-Def3b*), and PRPs (*Lser-PRP1* and *Lser-PRP4*) are only up-regulated upon exposure to *P. aeruginosa* (Figure 1; Table 1). *D. melanogaster edin* is also induced upon infection (Verleyen et al., 2006), and at least two homologs of *edin* are upregulated upon systemic infection in the house fly (Sackton et al., 2017). This suggests that Edin proteins are an important, and conserved, component of the immune response across flies, with bacteria-specific induction. All observations were expected given bacteria-induced expression and anti-bacterial activity of Edin and PRPs in *L. sericata* and other flies (Cytryńska et al., 2020; Gordon et al., 2008; Levashina et al., 1995; Vanha-Aho et al., 2012). Therefore, the *L. sericata* response to feeding on *P. aeruginosa* identifies multiple expected candidates for further study as biomimetic anti-microbials targeted toward specific pathogens, including canonical AMPs.

### Non-immune genes differentially expressed after exposure to A. baumannii

Genes without any direct relationships to immunity were upregulated upon exposure to *A. baumannii* and are suggestive of the signaling pathways involved in the response of *L. sericata* to bacterial exposure. Two induced non-immune genes are homologous to *D. melanogaster* genes (*uri* and *Ncoa6*) involved in signaling pathways and transcriptional regulation (Table 1). *Ncoa6* is a member of the Hippo signaling pathway, which regulates tissue growth (Qing et al., 2014), and *uri* is involved in nutrient signaling associated with the Insulin-like receptor (InR)/Target of rapamycin (TOR) pathway (Chaves-Pérez et al., 2018; Kirchner et al., 2008). When *uri* is not differentially expressed (i.e., in the *P. aeruginosa* treatment), *Lst*, which is also associated with InR/TOR signaling (Alfa et al., 2015), is upregulated (Supplementary Table S4). Upregulation of *ecd* was also observed after treatment with *A. baumanii* (Supplementary Table S5). In *D. melanogaster*, Ecd physically interacts with Uri (Rhee et al., 2014). Another differentially expressed gene (*FF38_01932*) is homologous to *D. melanogaster Edg84A* (Figure 5), which is involved in formation of the pupal cuticle (Apple and Fristrom, 1991). Expression of *Edg84A* in *D. melanogaster* is regulated by nuclear hormone receptor transcription factors (Akagi et al., 2013; Murata et al., 1996). Nuclear receptors respond to ecdysteroid pulses to trigger developmental transitions (Yamada et al., 2000; Yamanaka et al., 2013). These results are consistent with experiments in *D. melanogaster* finding immune responses impacting hormonal signaling pathways that regulate development (DiAngelo et al., 2009; Lee and Lee, 2018; Suzawa et al., 2019. Because induction of these non-immune genes is greater for *A. baumanii* exposure, we hypothesize that *A. baumanii* has greater effects on growth and developmental physiology in *L. sericata*.

While the non-immune responses were not expected, they are not surprising given that this experiment exposed flies to bacteria through feeding. When all non-immune genes discussed here are taken together a larger gut-oriented picture emerges (Liu et al. 2017, Wang et al. 2020) (Figure 6). *Peritrophin-15*, a component of the peritrophic matrix (Wijffels et al., 2001), is downregulated at 1 h after *P. aeruginosa* treatment (Supplementary Table S4). In addition, *Osiris* genes were suppressed after 1 h of feeding (Figure 6A). Osiris proteins are gut expressed, located in the cellular membrane, and possess chitin binding domains, implying a role in the peritrophic matrix (Dorer et al., 2003; Shah et al., 2012; Smith et al., 2018). Intriguingly, they are also differentially expressed when *Drosophila sechellia* feed on toxic octanoic acid (Lanno et al., 2017), and may be involved in adaptation of *D. sechellia* to tolerate that compound in their host plant, *Morinda citrifolia* (Andrade López et al., 2017; Lanno et al., 2019). In addition, three transcripts homologous to *D. melanogaster* genes encoding proteins containing chitin binding domains (*Edg84A* and two *Ccp84A* genes) are also downregulated by *A. baumanii* exposure (Figure 6A). Downregulation of these 7 transcripts encoding proteins likely to be present in the peritrophic matrix suggests changes to the gut epithelium in *L. sericata* and aligns with observations in *D. melanogaster* (Buchon et al., 2009).

**Figure 6.**
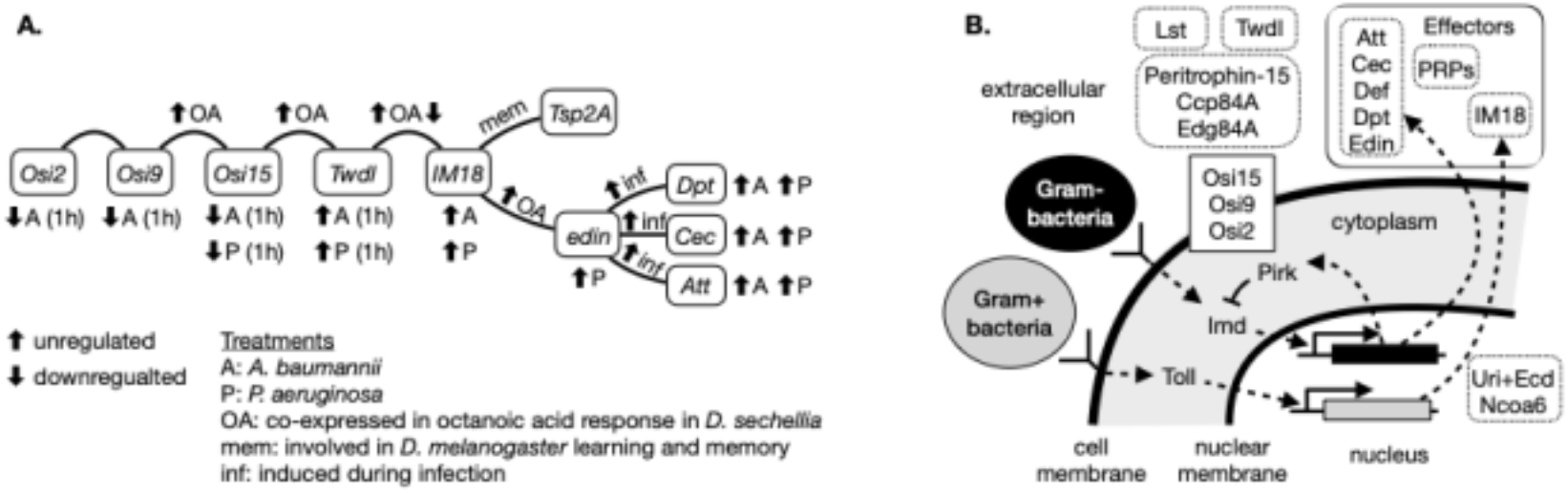
Genes induced in *L. sericata* by feeding on *P. aeruginosa* and *A. baumanii* are related through coexpression in other experiments, shared regulatory pathways, and known physical interactions. **A.** Genes that are differentially expressed in *L. sericata* upon exposure to *P. aeruginosa* (P) and *A. baumanii* (A), along with one gut expressed gene that is not differentially expressed (*Tsp2A*), are connected by curved lines based on co-expression in other experiments, shared functions, or membership in the same gene families. Other experiments include those in which *D. sechellia* was fed octanoic acid (OA), inferred functions in learning and memory in *D. melanogaster*, or induction during bacterial infection (Bozler et al., 2017; Gordon et al., 2008; Lanno et al., 2017; Short and Lazzaro, 2013). **B.** Annotated cellular localization of proteins encoded by genes differentially expressed after bacterial exposure in *L. sericata* are shown. The relationships of the differentially expressed genes to the two primary cell signaling pathways involved in the humoral innate immune response (Toll and Imd) are also shown, along with their (imperfectly) dogmatic relationship to Gram-negative (Gram-) and Gram-positive (Gram+) bacteria.

Other genes identified as differentially expressed in *D. sechellia* after exposure to octanoic acid were also differentially expressed in *L. sericata* after *A. baumanii* exposure, including *Twdl* genes, *IM18*, and *edin* (Lanno et al., 2017). *IM18* is also implicated in *D. melanogaster* learning and memory, along with *Tsp2A* (Bozler et al., 2017). *Tsp2A* is another gene that is expressed in the gut and is important for midgut development (Izumi et al., 2016; Xu et al., 2019). In our experiment and in the octanoic acid experiment in *D. sechellia, edin* and *IM18* are differentially expressed after gut exposure as a result of an external environmental stimulus and both are exported from cells (Figure 6B).

The induction of non-immune genes in the gut after feeding, including the *Osiris* and *Twdl* genes, suggests that either the Imd signaling pathway drives expression of non-immune genes or other signaling pathways are being induced in response to something other than infection. Nearly all of the immune genes that are induced in *L. sericata* by *P. aeruginosa* and *A. baumanii* (e.g., *edin*, *pirk*, and AMP genes) are regulated by the Imd pathway, which is induced by Gram-negative bacteria (Kleino et al., 2008; Kleino and Silverman, 2014; Vanha-Aho et al., 2012). However, *IM18* is considered to be regulated by Toll (Seto and Tamura, 2013; Uttenweiler-Joseph et al., 1998). The Toll pathway is regulated by Hippo signaling (Liu et al., 2016), which regulates the proliferation of gut stem cells, including after injury (Ren et al., 2010). One of the genes that is differentially expressed by *A. baumanii* exposure (*Ncoa6*) is itself a regulator of Hippo signaling (Qing et al., 2014), suggesting that the Toll pathway is indeed activated when *L. sericata* eat Gram-negative bacteria.

One explanation for the induction of non-immune genes when *L. sericata* is exposed to *A. baumanii* is that the larval gut is responding to tissue damage. The induction of AMP and effector genes was less when *L. sericata* was exposed to *A. baumanii* relative to *P. aeruginosa* (Figure 1; Figure 2). The weaker immune response raises the possibility that *A. baumanii* proliferates in the gut. The induction of multiple non-immune genes related to Toll/Hippo signaling and wound healing (including *Ncoa6* and *uri/ecd*) by exposure to *A. baumanii* (Table 1) suggest damage to the gut epithelium that must be repaired (Ren et al., 2010). The downregulation of genes expected to encode components of the peritrophic matrix (*Peritrophin 15*, *Edge84A*, *Osiris* and *Ccp84A* genes) could be relatedto gut damage—the peritrophic matrix plays an important role in reducing the harmful effects of bacterial infection in the fly gut (Kuraishi et al., 2011; Weiss et al., 2014).

Our results are consistent with other evidence suggesting a more complex relationship between Imd and Toll pathways beyond merely responding to two different classes of microbial pathogens (Nishide et al., 2019; Valanne et al., 2011). Across the analyses conducted in these experiments, a broader picture emerges of interactions between infection and tissue damage mediated by Toll, Imd, InR/TOR, Hippo, and ecdysone signaling. Several of the same non-immune genes are differentially expressed in response to bacteria or chemical toxins in the gut in *Drosophila* and *Lucilia*, with *IM18* and *edin* potentially linking the Imd and Toll pathways in the gut, and the Toll pathway potentially functioning in response to gut damage.

### Conclusions

In summary, we demonstrate induction of AMP and other effector genes as the primary differential expression occurring when *L. sericata* second instar larvae are exposed, by feeding, to *P. aeruginosa* and *A. baumannii. P. aeruginosa* induced more AMPs, and the induction of effectors increases over time with exposure to *P. aeruginosa*, but not for *A. baumanii. L. sericata* exhibited distinct responses to these bacteria and the similarities and differences in the differential expression associated with them helps to dissect the role of the gut and genes in these taxon-specific responses.

## Acknowledgements

The authors would like to thank Ashleigh Faris, Jonathan Parrot, Elida Espinoza, and Meaghan Pimsler at Texas A&M, Rick Metz and Charlie Johnson at the Texas A&M AgriLife Genomics and Bioinformatics Service, and Kranti Konganti at the Texas A&M Institute for Genome Sciences and Society for their assistance on this project. This work was completed in part using the Maxwell Cluster provided by the Research Computing Data Core at the University of Houston and the High Performance Computing Cluster at the Texas A&M Institute for Genome Sciences and Society. Josh Benoit at the University of Cincinnati and Sheena Cotter at the University of Lincoln provided helpful commentary.

## References cited

Akagi K, Kageyama Y, Kayashima Y, et al. (2013) The binding of multiple nuclear receptors to a single regulatory region is important for the proper expression of EDG84A in *Drosophila melanogaster*. Journal of Molecular Biology 425(1): 71–81.

Alfa RW, Park S, Skelly K-R, et al. (2015) Suppression of insulin production and secretion by a decretin hormone. Cell Metabolism 21(2): 323–334.

Andersen AS, Joergensen B, Bjarnsholt T, et al. (2010) Quorum-sensing-regulated virulence factors in *Pseudomonas aeruginosa* are toxic to *Lucilia sericata* maggots. Microbiology 156(Pt 2): 400–407.

Anderson GS, Byrd JH and Castner JL (2001) Insect succession on carrion and its relationship to determining time of death. Forensic entomology: the utility of arthropods in legal investigations 143. CRC Press Boca Raton, FL, USA: 76.

Andrade López JM, Lanno SM, Auerbach JM, et al. (2017) Genetic basis of octanoic acid resistance in *Drosophila sechellia:* functional analysis of a fine-mapped region. Molecular Ecology 26(4): 1148–1160.

Anstead CA, Korhonen PK, Young ND, et al. (2015) *Lucilia cuprina* genome unlocks parasitic fly biology to underpin future interventions. Nature Communications 6: 7344.

Apple RT and Fristrom JW (1991) 20-Hydroxyecdysone is required for, and negatively regulates, transcription of *Drosophila* pupal cuticle protein genes. Developmental Biology 146(2): 569–582.

Barnes KM, Gennard DE and Dixon RA (2010) An assessment of the antibacterial activity in larval excretion/secretion of four species of insects recorded in association with corpses, using *Lucilia sericata* Meigen as the marker species. Bulletin of Entomological Research 100(6): 635–640.

Benjamini Y and Hochberg Y (1995) Controlling the false discovery rate: a practical and powerful approach to multiple testing. Journal of the Royal Statistical Society. Series B, Statistical Methodology 57(1). [Royal Statistical Society, Wiley]: 289–300.

Benoit JB, Vigneron A, Broderick NA, et al. (2017) Symbiont-induced odorant binding proteins mediate insect host hematopoiesis. eLife 6. DOI: 10.7554/eLife.19535.

Bexfield A, Bond AE, Roberts EC, et al. (2008) The antibacterial activity against MRSA strains and other bacteria of a <500Da fraction from maggot excretions/secretions of *Lucilia sericata* (Diptera: Calliphoridae). Microbes and Infection / Institut Pasteur 10(4):25–333.

Bianchi F, Flisi S, Careri M, et al. (2019) Vertebrate odorant binding proteins as antimicrobial humoral components of innate immunity for pathogenic microorganisms. PLoS One 14(3): e0213545.

Boucher HW, Talbot GH, Bradley JS, et al. (2009) Bad bugs, no drugs: no ESKAPE! An update from the Infectious Diseases Society of America. Clinical infectious diseases: an official publication of the Infectious Diseases Society of America 48(1): 1–12.

Bozler J, Kacsoh BZ, Chen H, et al. (2017) A systems level approach to temporal expression dynamics in *Drosophila* reveals clusters of long term memory genes. PLoS Genetics 13(10): e1007054.

Bray NL, Pimentel H, Melsted P, et al. (2016) Near-optimal probabilistic RNA-seq quantification. Nature Biotechnology 34(5): 525–527.

Buchon N, Broderick NA, Poidevin M, et al. (2009) *Drosophila* intestinal response to bacterial infection: activation of host defense and stem cell proliferation. Cell Host & Microbe 5(2): 200–211.

Buchon N, Silverman N and Cherry S (2014) Immunity in *Drosophila melanogaster--from* microbial recognition to whole-organism physiology. Nature Reviews Immunology 14(12): 796–810.

Bulet P, Hetru C, Dimarcq JL, et al. (1999) Antimicrobial peptides in insects; structure and function. Developmental and Comparative Immunology 23(4-5): 329–344.

Cazander G, van Veen KEB, Bouwman LH, et al. (2009) The influence of maggot excretions on PAO1 biofilm formation on different biomaterials. Clinical Orthopaedics and Related Research 467(2): 536–545.

CDC (2020) Antibiotic Resistance Threatens Everyone. Available at: https://www.cdc.gov/drugresistance/index.html (accessed 15 July 2020).

Chaves-Pérez A, Thompson S and Djouder N (2018) Roles and Functions of the Unconventional Prefoldin URI. In: Djouder N (ed.) Prefoldins: The New Chaperones. Cham: Springer International Publishing, pp. 95–108.

Chopra I, Schofield C, Everett M, et al. (2008) Treatment of health-care-associated infections caused by Gram-negative bacteria: a consensus statement. The Lancet Infectious Diseases 8(2): 133–139.

Cytryńska M, Rahnamaeian M, Zdybicka-Barabas A, et al. (2020) Proline-rich antimicrobial peptides in medicinal maggots of *Lucilia sericata* interact with bacterial *DnaK* but do not inhibit protein synthesis. Frontiers in Pharmacology 11: 532.

DiAngelo JR, Bland ML, Bambina S, et al. (2009) The immune response attenuates growth and nutrient storage in *Drosophila* by reducing insulin signaling. Proceedings of the National Academy of Sciences of the United States of America 106(49): 20853–20858.

Dorer DR, Rudnick JA, Moriyama EN, et al. (2003) A family of genes clustered at the Triplo-lethal locus of *Drosophila melanogaster* has an unusual evolutionary history and significant synteny with *Anopheles gambiae*. Genetics 165(2): 613–621.

Dubey SK and Tapadia MG (2018) *Yorkie* regulates neurodegeneration through canonical pathway and innate immune response. Molecular Neurobiology 55(2): 1193–1207.

Galardi S, Fatica A, Bachi A, et al. (2002) Purified box C/D snoRNPs are able to reproduce site-specific 2’-O-methylation of target RNA *in vitro*. Molecular and cellular biology 22(19): 6663–6668.

Glass CK and Ogawa S (2006) Combinatorial roles of nuclear receptors in inflammation and immunity. Nature Reviews Immunology 6(1): 44–55.

Gordon MD, Ayres JS, Schneider DS, et al. (2008) Pathogenesis of *Listeria-infected Drosophila wntD* mutants is associated with elevated levels of the novel immunity gene *edin*. PLoS pathogens 4(7): e1000111.

Hanson MA, Dostálová A, Ceroni C, et al. (2019) Synergy and remarkable specificity of antimicrobial peptides *in vivo* using a systematic knockout approach. eLife 8. DOI: 10.7554/eLife.44341.

Hirsch R, Wiesner J, Marker A, et al. (2019) Profiling antimicrobial peptides from the medical maggot *Lucilia sericata* as potential antibiotics for MDR Gram-negative bacteria. The Journal of Antimicrobial Chemotherapy 74(1): 96–107.

Hoffmann JA (2003) The immune response of *Drosophila*. Nature 426(6962): 33–38.

Hong L, Li X, Zhou D, et al. (2018) Role of *Hippo* signaling in regulating immunity. Cellular & Molecular Immunology 15(12): 1003–1009.

Huang Z-P, Zhou H, He H-L, et al. (2005) Genome-wide analyses of two families of snoRNA genes from *Drosophila melanogaster*, demonstrating the extensive utilization of introns for coding of snoRNAs. RNA 11(8): 1303–1316.

Imler J-L and Bulet P (2005) Antimicrobial peptides in *Drosophila:* structures, activities and gene regulation. Chemical Immunology and Allergy 86: 1–21.

Izumi Y, Motoishi M, Furuse K, et al. (2016) A tetraspanin regulates septate junction formation in *Drosophila* midgut. Journal of Cell Science 129(6): 1155–1164.

Kerridge A, Lappin-Scott H and Stevens JR (2005) Antibacterial properties of larval secretions of the blowfly, *Lucilia sericata*. Medical and Veterinary Entomology 19(3): 333–337.

Khalil S, Jacobson E, Chambers MC, et al. (2015) Systemic bacterial infection and immune defense phenotypes in *Drosophila melanogaster*. Journal of Visualized Experiments: JoVE (99): e52613.

Kim D, Pertea G, Trapnell C, et al. (2013) TopHat2: accurate alignment of transcriptomes in the presence of insertions, deletions and gene fusions. Genome Biology 14(4): R36.

Kirchner J, Vissi E, Gross S, et al. (2008) *Drosophila Uri*, a PP1alpha binding protein, is essential for viability, maintenance of DNA integrity and normal transcriptional activity. BMC Molecular Biology 9: 36.

Kleino A and Silverman N (2014) The *Drosophila* IMD pathway in the activation of the humoral immune response. Developmental and Comparative Immunology 42(1): 25–35.

Kleino A, Myllymäki H, Kallio J, et al. (2008) *Pirk* is a negative regulator of the *Drosophila* Imd pathway. Journal of Immunology 180(8): 5413–5422.

Kuraishi T, Binggeli O, Opota O, et al. (2011) Genetic evidence for a protective role of the peritrophic matrix against intestinal bacterial infection in *Drosophila melanogaster*. Proceedings of the National Academy of Sciences of the United States of America 108(38): 15966–15971.

Lanno SM, Gregory SM, Shimshak SJ, et al. (2017) Transcriptomic analysis of octanoic acid response in *Drosophila sechellia* using RNA-sequencing. G3 7(12): 3867–3873.

Lanno SM, Shimshak SJ, Peyser RD, et al. (2019) Investigating the role of Osiris genes in *Drosophila sechellia* larval resistance to a host plant toxin. Ecology and Evolution 9(4): 1922–1933.

Lee K-A and Lee W-J (2018) Immune–metabolic interactions during systemic and enteric infection in *Drosophila*. Current Opinion in Insect Science 29: 21–26.

Lemaitre B and Hoffmann J (2007) The host defense of *Drosophila melanogaster*. Annual Review of Immunology 25: 697–743.

Levashina EA, Ohresser S, Bulet P, et al. (1995) *Metchnikowin*, a novel immune-inducible proline-rich peptide from *Drosophila* with antibacterial and antifungal properties. European journal of biochemistry / FEBS 233(2). Wiley Online Library: 694–700.

Levy F, Bulet P and Ehret-Sabatier L (2004) Proteomic analysis of the systemic immune response of *Drosophila*. Molecular & Cellular Proteomics: MCP 3(2): 156–166.

Linger RJ, Belikoff EJ, Yan Y, et al. (2016) Towards next generation maggot debridement therapy: transgenic *Lucilia sericata* larvae that produce and secrete a human growth factor. BMC Biotechnology 16: 30.

Liu B, Zheng Y, Yin F, et al. (2016) Toll receptor-mediated *Hippo* signaling controls innate immunity in *Drosophila*. Cell 164(3): 406–419.

Liu X, Hodgson JJ, Buchon N (2017) *Drosophila* as a model for homeostatic, antibacterial, and antiviral mechanisms in the gut. PLoS Pathogens 13(5): e1006277.

Lynham J and Houry WA (2018) The multiple runctions of the PAQosome: An R2TP- and URI1 prefoldin-based chaperone complex. In: Djouder N (ed.) Prefoldins: The New Chaperones. Cham: Springer International Publishing, pp. 37–72.

McCarthy DJ, Chen Y and Smyth GK (2012) Differential expression analysis of multifactor RNA-Seq experiments with respect to biological variation. Nucleic Acids Research 40(10): 4288–4297.

Michel T, Reichhart JM, Hoffmann JA, et al. (2001) Drosophila Toll is activated by Gram-positive bacteria through a circulating peptidoglycan recognition protein. Nature 414(6865): 756–759.

Morgulis A, Coulouris G, Raytselis Y, et al. (2008) Database indexing for production MegaBLAST searches. Bioinformatics 24(16): 1757–1764.

Murata T, Kageyama Y, Hirose S, et al. (1996) Regulation of the *EDG84A* gene by *FTZ-F1* during metamorphosis in *Drosophila melanogaster*. Molecular and Cellular Biology 16(11): 6509–6515.

Nishide Y, Kageyama D, Yokoi K, et al. (2019) Functional crosstalk across IMD and Toll pathways: insight into the evolution of incomplete immune cascades. Proceedings. Biological sciences / The Royal Society 286(1897): 20182207.

Pavillard ER and Wright EA (1957) An antibiotic from maggots. Nature 180(4592): 916–917.

Pechal JL, Benbow ME, Crippen TL, et al. (2014) Delayed insect access alters carrion decomposition and necrophagous insect community assembly. Ecosphere 5(4): art45.

Pei Z, Sun X, Tang Y, et al. (2014) Cloning, expression, and purification of a new antimicrobial peptide gene from *Musca domestica* larva. Gene 549(1): 41–45.

Pelosi P, Iovinella I, Zhu J, et al. (2018) Beyond chemoreception: diverse tasks of soluble olfactory proteins in insects. Biological reviews of the Cambridge Philosophical Society 93(1): 184–200.

Pöppel A-K, Vogel H, Wiesner J, et al. (2015) Antimicrobial peptides expressed in medicinal maggots of the blow fly *Lucilia sericata* show combinatorial activity against bacteria. Antimicrobial Agents and Chemotherapy 59(5): 2508–2514.

Qing Y, Yin F, Wang W, et al. (2014) The Hippo effector *Yorkie* activates transcription by interacting with a histone methyltransferase complex through *Ncoa6*. eLife 3. DOI: 10.7554/eLife.02564.

Ren F, Wang B, Yue T, et al. (2010) Hippo signaling regulates *Drosophila* intestine stem cell proliferation through multiple pathways. Proceedings of the National Academy of Sciences of the United States of America 107(49): 21064–21069.

Rhee DY, Cho D-Y, Zhai B, et al. (2014) Transcription factor networks in *Drosophila melanogaster*. Cell reports 8(6): 2031–2043.

Ritchie ME, Phipson B, Wu D, et al. (2015) limma powers differential expression analyses for RNA-sequencing and microarray studies. Nucleic Acids Research 43(7): e47.

Robinson MD, McCarthy DJ and Smyth GK (2010) edgeR: a Bioconductor package for differential expression analysis of digital gene expression data. Bioinformatics 26(1): 139–140.

Sackton TB, Lazzaro BP, Schlenke TA, et al. (2007) Dynamic evolution of the innate immune system in *Drosophila*. Nature genetics 39(12): 1461–1468.

Sackton TB, Lazzaro BP and Clark AG (2017) Rapid expansion of immune-related gene families in the house fly, *Musca domestica*. Molecular Biology and Evolution 34(4): 857–872.

Scocchi M, Tossi A and Gennaro R (2011) Proline-rich antimicrobial peptides: converging to a non-lytic mechanism of action. Cellular and Molecular Life Sciences: CMLS 68(13): 2317–2330.

Seto Y and Tamura K (2013) Extensive differences in antifungal immune response in two *Drosophila* species revealed by comparative transcriptome analysis. International Journal of Genomics and Proteomics 2013: 542139.

Shah N, Dorer DR, Moriyama EN, et al. (2012) Evolution of a large, conserved, and syntenic gene family in insects. G3 2(2): 313–319.

Sherman RA (2014) Mechanisms of maggot-induced wound healing: what do we know, and where do we go from here? Evidence-based complementary and alternative medicine: eCAM 2014: 592419.

Sherman RA, Hall MJ and Thomas S (2000) Medicinal maggots: an ancient remedy for some contemporary afflictions. Annual Review of Entomology 45: 55–81.

Sherman RA, Mumcuoglu KY, Grassberger M, et al. (2013) Maggot therapy. In: Grassberger M, Sherman RA, Gileva OS, et al. (eds) Biotherapy - History, Principles and Practice: A Practical Guide to the Diagnosis and Treatment of Disease Using Living Organisms. Dordrecht: Springer Netherlands, pp. 5–29.

Short SM and Lazzaro BP (2013) Reproductive status alters transcriptomic response to infection in female *Drosophila melanogaster*. G3 3(5): 827–840.

Sigrist CJA, Cerutti L, de Castro E, et al. (2010) PROSITE, a protein domain database for functional characterization and annotation. Nucleic acids research 38(Database issue): D161–6.

Smith CR, Morandin C, Noureddine M, et al. (2018) Conserved roles of Osiris genes in insect development, polymorphism and protection. Journal of Evolutionary Biology 31(4): 516–529.

