## Supplemental Material for "Gene expression in *Lucilia sericata* (Diptera: Calliphoridae) larvae exposed to *Pseudomonas aeruginosa* and *Acinetobacter baumanii* identifies shared and microbe-specific induction of immune genes"

### Supplemental Figures

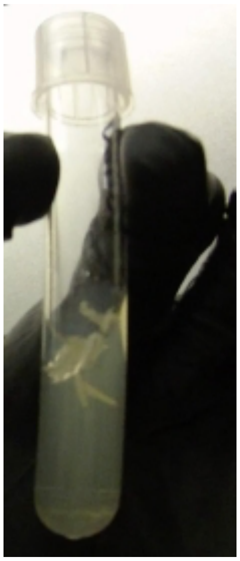

**Supplemental Figure S1**. Larvae on and just below surface of agar in 5 mL round bottom tube.

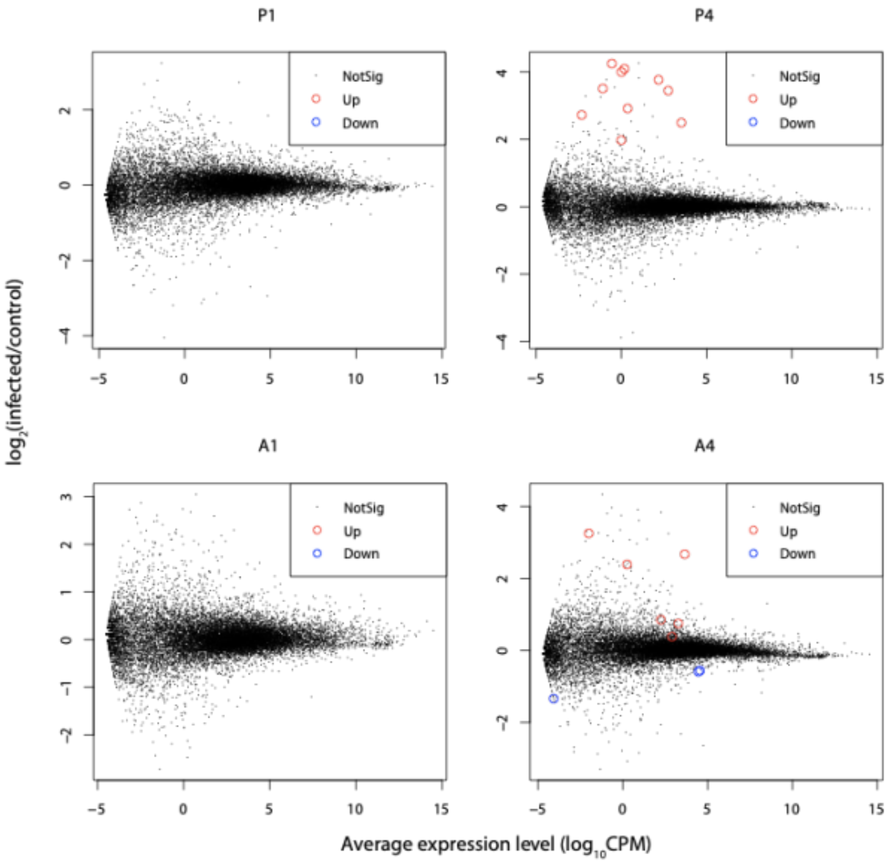


**Supplemental Figure S2.** The global patterns of differential expression across microbial treatments. X-axis indicates the average $\log_{10}$ counts per million (CPM). Y-axis indicates $\log_2$ of the ratio of gene expression level of infected (bacterial feeding) relative to control. P (*Pseudomonas*) and A (*Acinetobacter*) indicate the bacterial treatment. 1 and 4 indicate the duration of the feeding treatment (in hours). Upregulated and down regulated genes are shown as red and blue dots respectively. The rest of the genes are shown as black dots. Data are from kallisto alignment to the i5k genome, with differential expression analysis performed using edgeR/limma. All results can be found in Supplementary Tables S4-S5.

### Supplemental Tables

**Supplemental Table S1.** Sample information.

| Treatment | Exposure time (h) | Replicate | Batch 1 IDs | Batch 2 IDs | Batch 3 IDs |
|---|---|---|---|---|---|
| **PBS (control)** | 1 | A | C1A1 | C1A3 | C1A4 |
| | | B | C1B1 | C1B3 | C1B4 |
| | | C | C1C1 | C1C3 | C1C4 |
| | | D | C1D1 | C1D3 | C1D4 |
| ***P. aeruginosa*** | 1 | A | PA1A1 | PA1A3 | PA1A4 |
| | | B | PA1B1 | PA1B3 | PA1B4 |
| | | C | PA1C1 | PA1C3 | PA1C4 |
| | | D | PA1D1 | PA1D3 | PA1D4 |
| ***A. baumannii*** | 1 | A | AB1A1 | AB1A3 | AB1A4 |
| | | B | AB1B1 | AB1B3 | AB1B4 |
| | | C | AB1C1 | AB1C3 | AB1C4 |
| | | D | AB1D1 | AB1D3 | AB1D4 |
| **PBS (control)** | 4 | A | C4A1 | C4A3 | C4A4 |
| | | B | C4B1 | C4B3 | C4B4 |
| | | C | C4C1 | C4C3 | C4C4 |
| | | D | C4D1 | C4D3 | C4D4 |
| ***P. aeruginosa*** | 4 | A | PA4A1 | PA4A3 | PA4A4 |
| | | B | PA4B1 | PA4B3 | PA4B4 |
| | | C | PA4C1 | PA4C3 | PA4C4 |
| | | D | PA4D1 | PA4D3 | PA4D4 |
| ***A. baumannii*** | 4 | A | AB4A1 | AB4A3 | AB4A4 |
| | | B | AB4B1 | AB4B3 | AB4B4 |
| | | C | AB4C1 | AB4C3 | AB4C4 |
| | | D | AB4D1 | AB4D3 | AB4D4 |

**Supplemental Table S2.** Differential expression results for exposure to *Pseudomonas aeruginosa* using the model Expression = Treatment + hours (E~T+h), as well as the model with an interaction term (E~T+h+T×h). RNA-seq reads were aligned to the *L. sericata* transcriptome using kallisto.

**Supplemental Table S3.** Differential expression results for exposure to *Acinetobacter baumannii* using the model Expression = Treatment + hours (E~T+h), as well as the model with an interaction term (E~T+h+T×h). RNA-seq reads were aligned to the *L. sericata* transcriptome using kallisto.

**Supplemental Table S4.** Differential expression results for comparisons within timepoints for *Pseudomonas* treatment (E~T). RNA-seq reads were aligned to either the Melbourne or i5k *L. cuprina* assembly and annotation, and differential expression analysis was performed using either kallisto/limma/edgeR or TopHat2/cuffdiff.

**Supplemental Table S5.** Differential expression results for comparisons within timepoints for *Acinetobacter* treatment (E~T). RNA-seq reads were aligned to either the Melbourne or i5k *L. cuprina* assembly and annotation, and differential expression analysis was performed using either kallisto/limma/edgeR or TopHat2/cuffdiff.

**Supplemental Table S6.** Differential expression results for exposure to *Pseudomonas* using both models (E~T+h and E~T+h+T×h) and the Melbourne and i5k reference genomes with the kallisto/limma/edgeR analysis.

**Supplemental Table S7.** Differential expression results for exposure to *Acinetobacter* using both models (E~T+h and E~T+h+T×h) and the Melbourne and i5k reference genomes with the kallisto/limma/edgeR analysis.

**Supplemental Table S8.** Counts per million (CPM) values aligning reads from all treatments and controls to the *L. sericata* transcriptome with kallisto.

**Supplemental Table S9.** Counts per million (CPM) values aligning reads from all treatments and controls to the *L. cuprina* i5k assembly and annotation with kallisto.

**Supplemental Table S10.** Counts per million (CPM) values aligning reads from all treatments and controls to the *L. cuprina* Melbourne assembly and annotation with kallisto.
